# The ecomorphology of southern African rodent incisors: Potential applications to the hominin fossil record

**DOI:** 10.1101/429365

**Authors:** Oliver C.C. Paine, Jennifer N. Leichliter, Nico Avenant, Daryl Codron, Austin Lawrence, Matt Sponheimer

## Abstract

The taxonomic identification of mammalian fauna within fossil assemblages is a well-established component of paleoenvironmental reconstructions. However, many fragmentary specimens recovered from fossil sites are often disregarded as they can be difficult to identify with the precision required for taxonomic methods. For this reason, the large numbers of isolated rodent incisors that are often recovered from hominin fossil bearing sites have generally been seen as offering little interpretive value. Ecomorphological analysis, often referred to as a “taxon-free” method, can potentially circumvent this problem by focusing on the adaptive, rather than the taxonomic significance of rodent incisor morphology. Here, we determine if the morphology of the upper incisors of modern southern African rodents reflects dietary behavior using discriminant function analysis. Our model suggests that a strong ecomorphological signal exists in our modern sample and we apply these results to two samples of isolated incisors from the hominin fossil bearing sites, Sterkfontein and Swartkrans.

## Introduction

Fossil assemblages often contain elements that are of little or no interpretive value for traditional paleoecological investigations. In particular, methods relying on precise taxonomic identification of fossil specimens can rarely use fragmentary, or otherwise undiagnostic material. Such is the case with most rodent-based reconstructions of hominin paleohabitats, which rely on the preservation of the molar tooth rows for species, genus, and/or subfamily-level identification [e.g., 1–6]. While such material exists, in many fossil localities there is often an abundance of isolated incisors that heretofore have been of little interpretive value due to their poor taxonomic control. Because rodent incisors are ever-growing, they are not rooted in the alveoli in the same manner as other teeth. As such, rodent incisors often freely slide in and out of the maxillary and premaxillary alveoli when soft tissue is lost after death—a major reason why so many are found in isolation in fossil deposits—and in the course of our (OP and JL) work with fossil rodent collections from South African cave deposits we have come across an abundance of isolated incisors. The main objective of this study is to determine if these abundant fossil elements can yield an ecomorphological signal to augment previous interpretations of hominin paleohabitats.

Ecomorphological analysis is well established in paleontological research and has been successfully applied to reconstructions of hominin paleohabitats [e.g., 7–9]. These analyses are predicated on the fact that morphology often reflects functional adaptations to the ecological niche that a species occupies. Additionally, ecomorphological studies rest on the assumption that convergent morphological adaptations can arise across broadly related taxa (extant or extinct) in response to similar ecological pressures [10–13]. Thus, by understanding how specific adaptations in modern organisms correlate to ecological parameters, such as diet, locomotion, and habitat preference, inferences can be made about fossil specimens to improve paleoecological interpretations [see 14 for discussion].

For example, paleoanthropologists have used the limb morphology of fossil bovids to reconstruct hominin paleohabitats [7, 9, 14–17]. This has been particularly useful because bovid remains associated with fossil hominins, while generally abundant, can often be fragmentary and difficult to identify to precise taxonomic levels. Ecomorphological analyses circumvent this problem by focusing on the adaptive, rather than the taxonomic significance of morphological characteristics. Because of this, the method is often described as a “taxon-free” approach. That being said, phylogenetic proximity among species undeniably influences morphological correspondence in many instances [18–20] but often this is reflective of similar or shared adaptations as much as shared evolutionary history. Nonetheless, a major benefit of such taxon-free approaches is that they do not assume, as do most taxonomic methods, that fossil species had the same dietary and habitat tolerances as their closest living relatives, which we know is sometimes, if not often, unwarranted [e.g., 9, 15, 21, 22].

Small mammals (particularly rodents, though also shrews and sengis) can be well represented in hominin fossil assemblages and have increasingly been used to investigate early hominin habitats [e.g., 1–6, 23–26]. Because of their taxonomic diversity and relatively small home ranges, rodents arguably provide a more localized habitat signal compared to wider-ranging, large mammals in the fossil record. Additionally, because the majority of rodent species alive today are also found in the hominin fossil record [3], modern studies of rodent ecology can be directly applied to interpretations of the past.

In addition to being ever-growing, rodent incisors are curved, and lack enamel on their lingual surfaces. The exposed dentine wears more quickly than the enamel on the labial surface and thus allows the tooth to form a sharp, chisel-like structure with wear [see 27]. Because of this, the incisor morphology of any given rodent potentially represents an ecomorphological signal reflective of both long-term evolutionary adaptation and short-term behavioral proclivities.

Croft et al. [28] constructed a prediction model based on the morphology of rodent upper incisors that successfully assigned three broad dietary categories to extant South American species, and they argued that their model can potentially be applied to dietary interpretations of extinct taxa. Unfortunately, the success of the caviomorph model relies on a measurement (chord length) that we have never been able to assess on isolated fossil incisors. This measurement requires accurate location of the point on the enamel surface at which the incisor erupts from the premaxilla and, when using isolated incisors, there are obviously no instances of premaxillary association. What is more, after years of sorting and identifying rodent fossils from South African cave sites for this and related projects, we have yet to encounter an isolated incisor where this landmark is reliably preserved on the enamel surface.

Here, we contribute to the existing research on rodent incisor ecomorphology in two ways. First, we modify existing methods to determine if isolated incisors of southern African rodents can be used to provide dietary information. Second, we begin to assess the suitability of this method for use with the paleontological record by analyzing fossil rodent incisors from the South African hominin-bearing sites Sterkfontein (STS-4) and Swartkrans (SKX-1), which are temporally distinct and generally believed to represent different paleoenvironments [9, 29].

## Materials and methods

We sampled 41 modern South African rodent species curated in the collections of the Ditsong Museum of Natural History, Pretoria, South Africa (six Families and 31 Genera, *N*=163; S1 Table). For each species, we measured two adult males and two adult females, with the exception of the Cape mole-rat, *Georychus capensis*, for which reliable measurements could only be attained for three individuals. Specimens were photographed and measured at the museum using a Keyence^®^ VHX-2000E digital microscope with a VH-Z00R zoom lens (0x to 50x) and internal measurement software, and with Mitituyo^®^ Absolute AOS Digimatic calipers (for measuring mesiodistal diameter) (Fig 1).

**Fig 1.**
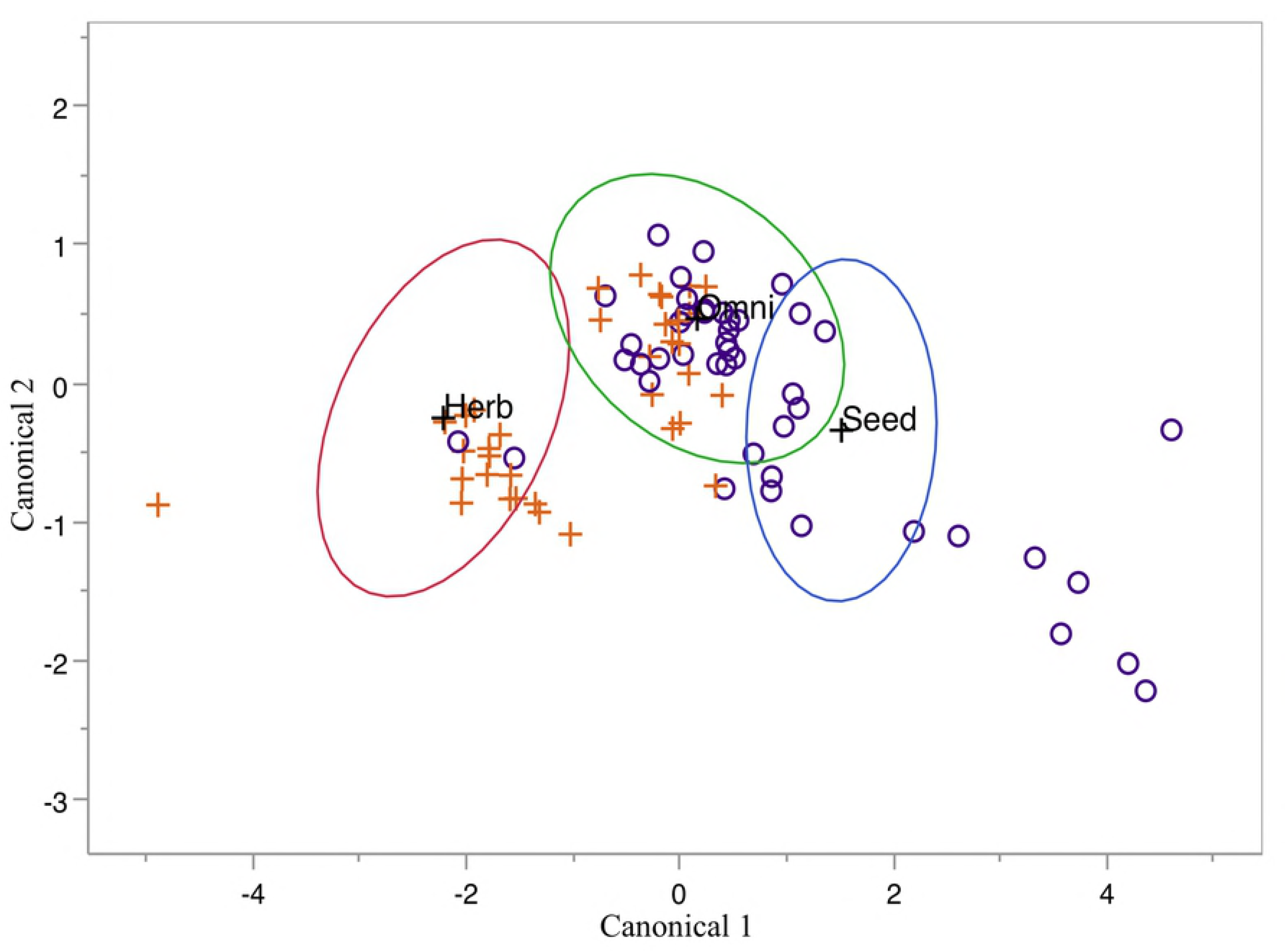
Measurements recorded. Upper incisors of *Rhabdomys pumilio* with the measurements adapted from Croft et al. [28] marked. Abbreviations: BD, buccolingual diameter; DE, distal enamel extent; MD, mesiodistal diameter; OD, buccolingual occlusal diameter; RC, outer radius of curvature; OA, occlusal angle. CL, chord length, was used by Croft et al. [28] but was unable to be measured on our fossil specimens (see text for explanation).

Prior to measurement, all specimens were mounted such that the distal surface of the left incisor faced upwards and was parallel to the microscope stage. Measurement protocol was adapted from Croft et al. [28]. Measurements were taken of buccolingual diameter (BD), distal enamel extent (DE), mesiodistal diameter (MD), buccolingual occlusal diameter (OD), and outer radius of curvature (RC) (Fig 1). Our values for DE represent the average of three enamel thickness measurements taken perpendicular to the superior, inferior, and midpoint of the lingual surface. We made this decision due to the difficulty in ascertaining the precise location of the enamel-dentine junction on some of the digital images. DE for taxa with grooved enamel surfaces (e.g., *Otomys sp*.) was measured in the same fashion, while acknowledging that the enamel structure of a grooved incisor is a more complex feature to quantify (Croft et al. [28] did not sample rodents with grooved incisors). To our knowledge, the functional significance of grooved incisors is unresolved and is potentially an artifact of benign genetic mutation [30].

We also recorded a new measurement, occlusal angle (OA), which is the angle created by the occlusal surface of the incisor and a radius drawn from the tooth’s inferior tip to the center of the circle inferred by its curvature (Fig 1). We chose this measurement as a means to measure the “sharpness” of the tooth’s cutting edge.

Similar to Croft et al. [28], three broad dietary categories were assigned: herbivore (predominantly leafy, green vegetable matter), seed eater (significant seed consumption, but also arthropods), and omnivore (vegetable matter, fruits, seeds, and arthropods). We include species that consume significant amounts of arthropods in our seed eater category because the hard, chitinous exoskeletons of many insects consumed (e.g., beetles) present a similar mechanical challenge. Also, we are aware that the term “omnivore” can have multiple definitions depending on the taxa being investigated and therefore “omnivore” lacks the precision one might prefer. However, it is not our goal to speak to those debates here and we use the term as convenient shorthand for the rodent diets noted above.

Dietary categories were assessed using Skinner and Chimimba [31] and Happold [32]. The dietary information for the species in our study was generally consistent between the two sources; however, when there were conflicting accounts, we opted to choose the description that referenced the most field studies and provided the most detail.

After measurements were taken, we calculated ratios using all possible combinations of linear measurements. We did this to better quantify the shape of the teeth and to mitigate the inherent size differences among our samples which range from the ∼6 g pygmy mouse (*Mus minutoides*) to the ∼3 kg springhare (*Pedetes capensis*). The median values for each species were used to calculate a *K* statistic [33] for each linear measurement and ratio in order to assess the strength of phylogenetic signal based on the phylogeny constructed by Faurby & Svenning [34]. *K* values > 1 suggest that phylogeny is driving the similarities between closely related species whereas values < 1 indicate that similarities between closely related species are less than would be expected under a Brownian motion model of trait evolution.

A stepwise quadratic discriminant analysis (QDA) with leave-one-out cross validation (“jackknifing”) was performed using measurements with *K* values < 1. Cross validation was also performed using a dataset of rodent specimens housed at The National Museum, Bloemfontein, South Africa. This validation set consisted of 24 specimens from 6 species: 4 seed eaters (*Dendromus mysticalis*); 4 herbivores (*Otomys irroratus); and 16 omnivores (4 specimens each from Micaelamys namaquensis, Gerbilliscus leucogaster, Rhabdomys pumilio*, and *Mastomys coucha*).

All statistical analyses were performed in *R*.

## Results

Fourteen of the 36 measures tested for a phylogenetic signal produced *K* values < 1 and all 14 were ratio measurements (S2 Table). These ratio measurements were used in our stepwise QDA and a model was constructed (using OA/MD, OD/DE/ BD/MD, and RC/MD) that misclassified 37 of the 163 training specimens (22.7%; Fig 2 and Table 1).

**Fig 2.**
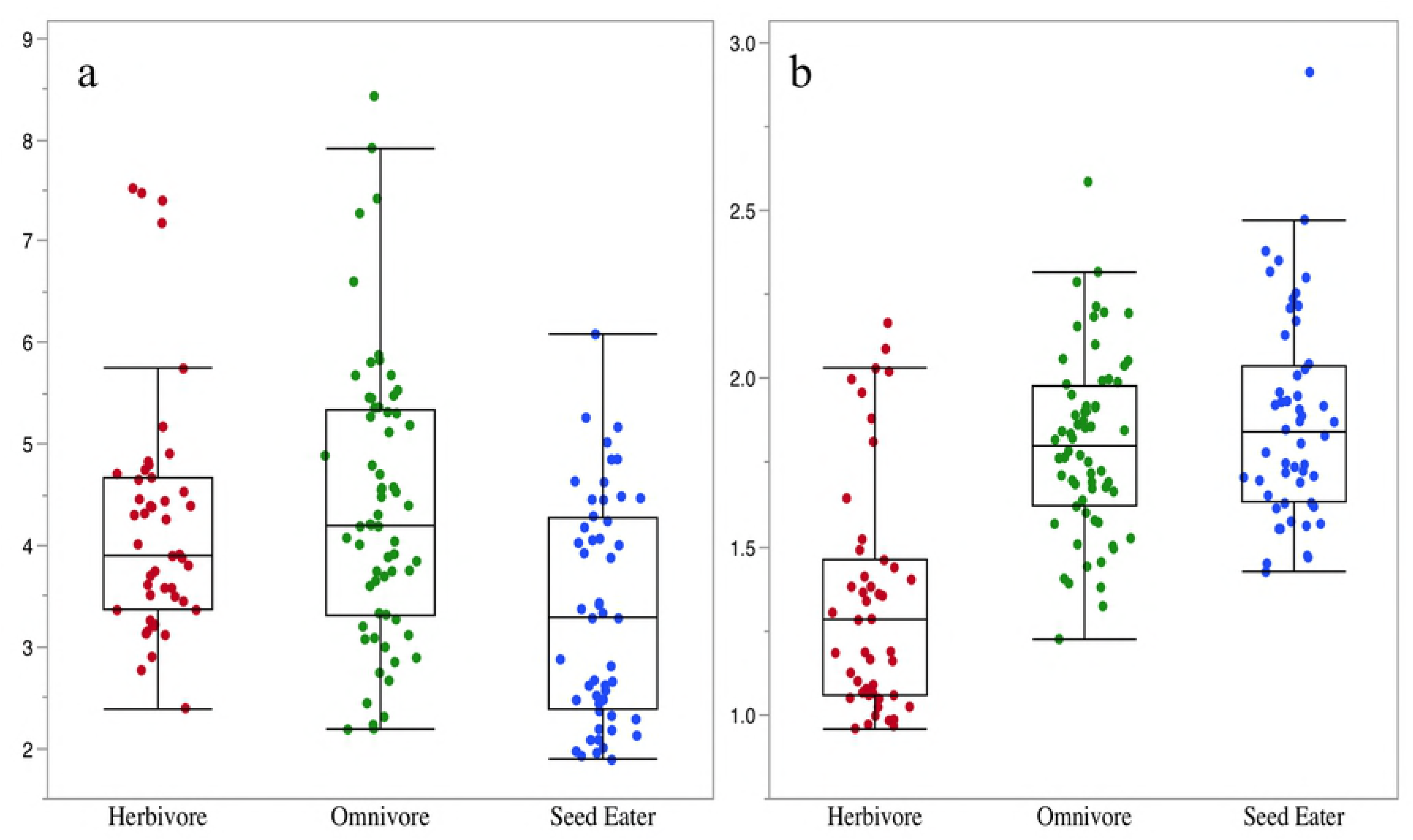
QDA model predicting modern rodent diets. The measures OA/MD, OD/DE, BD/MD, and RC/MD were used. Herbivores=red, omnivores=green, seed eaters=blue. Ellipses represent 50% of the distribution of each dietary category. Wilk’s Lambda=0.345, F=27.55, *P*< 0.0001.

**Table 1.** Predicted vs. actual diets of training specimens used in our QDA model.

|  | **Predicted Diet** |  |  |
| --- | --- | --- | --- |
| **Actual Diet** | Herbivore | Omnivore | Seed Eater |
| Herbivore | 34 | 12 | 1 |
| Omnivore | 4 | 49 | 11 |
| Seed Eater | 0 | 9 | 43 |

37 of 163 specimens misclassified (22.7%).

Canonical 1 explains 90% of the variance and is driven by BD/MD and OA/MD; Canonical 2 explains 10% of the variance and is driven by BD/MD and OD/DE (Table 2).

**Table 2.**
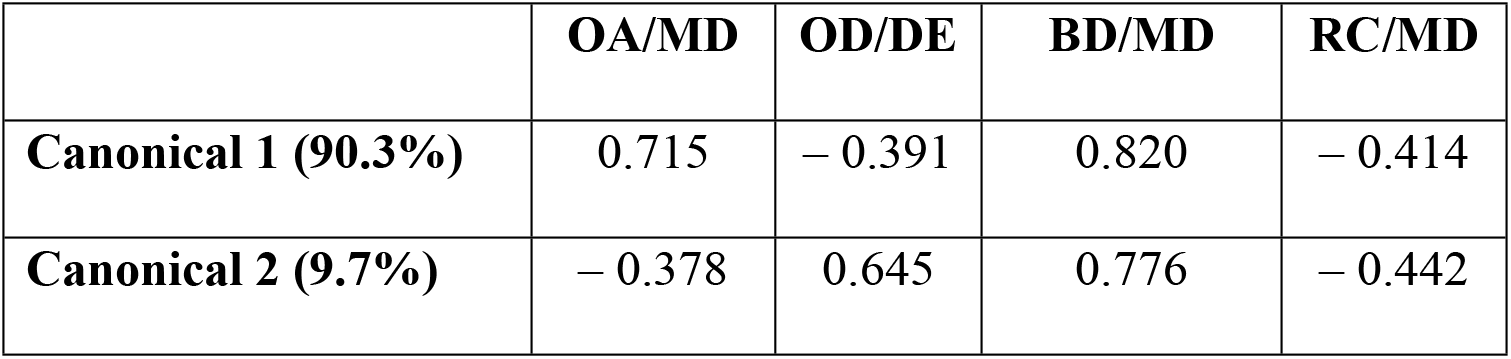
Standardized scoring coefficients of QDA model.

Twelve of the 47 herbivores were misclassified as omnivores and one was misclassified as a seed eater. Of the 64 omnivores in our training data, four were misclassified as herbivores and 11 as seed eaters. Nine of our 52 seed eaters were misclassified as omnivores and none were misclassified as herbivores (Table 1). Species for which at least three of the four specimens were misclassified included *Malacothrix typica* (4/4), *Mus indutus* (3/4), *Mus minutoides* (3/4), *Parotomys littledalei* (3/4), *Saccostomys capensis* (3/4), *Thallomys paedulcus*(4/4), and *Xerus inauris* (3/4) (S3 Table).

Leave-one-out cross validation produced a model that misclassified 41 of the 163 training specimens (25.1%), four more misclassifications than our original model. Our jackknifed model differed in its misclassifications of seed eaters, misclassifying all four *Desmodilus auricularis* specimens as omnivores, whereas the original model misclassified one. It also misclassified a single specimen of *Steatomys pratensis* as an omnivore whereas our original model correctly classified all four. When our validation specimens were entered into our QDA model, 4 of 24 (16.7%) were misclassified. All four seed eaters and all four herbivores were correctly classified whereas three omnivores were misclassified as herbivores (two specimens of *Micaelamys namaquensis* and one of *Rhabdomys pumilio*) and one omnivore specimen, *Mastomys coucha*, was misclassified as a seed eater.

## Discussion

### Modern rodent model

Our model successfully predicts the diets of African rodent taxa. The ratio data driving our dietary model suggest a discrete relationship between incisor morphology and the mechanical challenges posed by food items. In particular, the BD/MD and OD/DE ratios demonstrate that seed eaters have buccolingually deep incisors with a relatively thick enamel cap whereas herbivore incisors are mesiodistally broad and thin (Figs 3a and 3b).

**Fig 3a.**
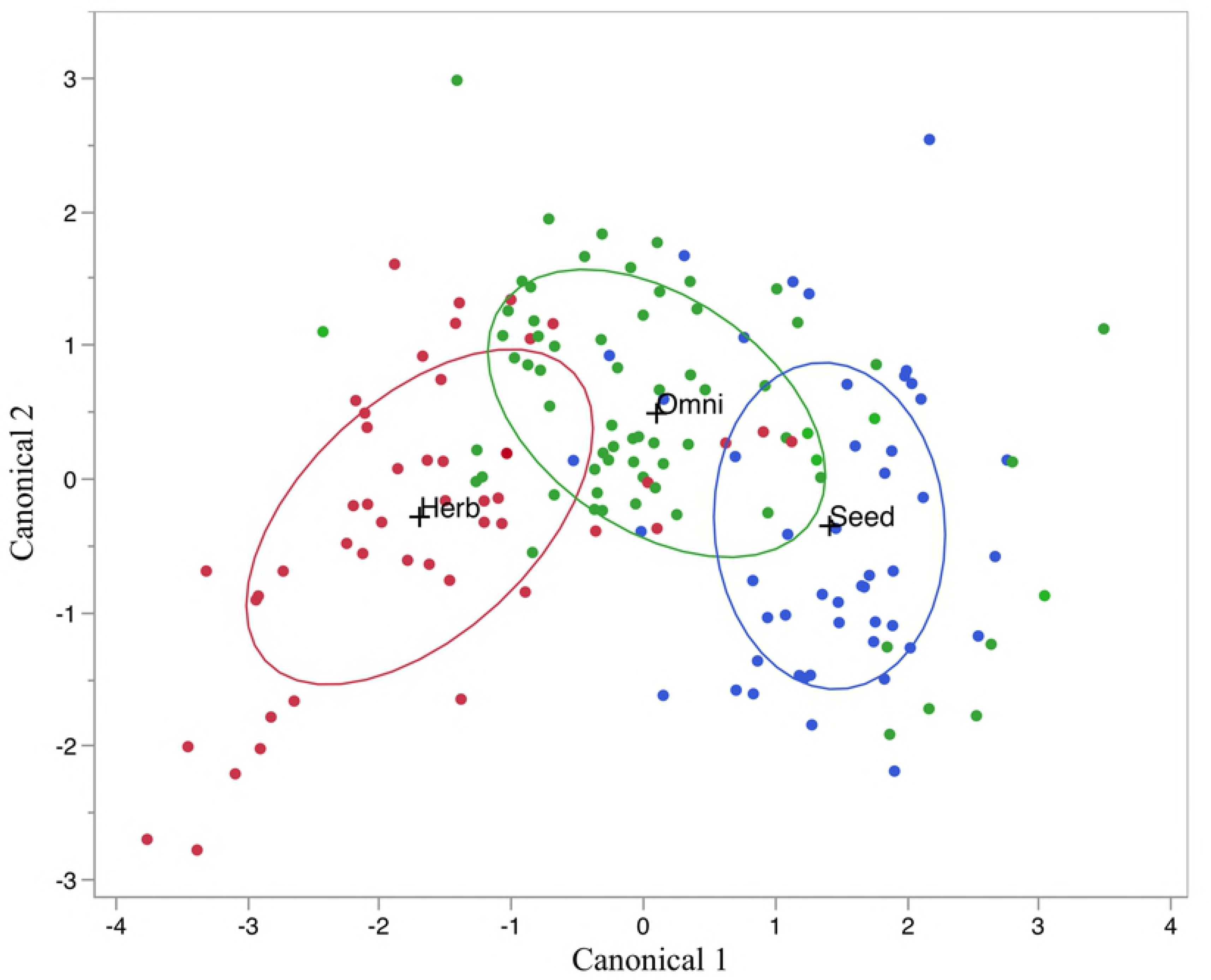
OD/DE of specimens used in the QDA model by dietary category. Seed eaters have thicker enamel caps relative to their occlusal surfaces. Kruskal-Wallis, *P*< 0.0001. Steel-Dwass all pairs comparisons: Herbivore-Omnivore, *P*= 0.6697; Herbivore-Seed Eater, *P* = 0.0025, Omnivore-Seed Eater, *P*= 0.0001. **Fig 3b. BD/MD of training specimens by dietary category.** Herbivores have relatively broader incisors. Kruskal-Wallis, *P*< 0.0001. Steel-Dwass all pairs comparisons: Herbivore-Omnivore, *P < 0.0001; Herbivore-Seed Eater, P* < 0.0001, Omnivore-Seed Eater, *P*= 0.5249.

The morphological dichotomy between our seed eaters and herbivores is in line with the findings of Croft et al. [28] whose “fruit-seed” and “grass-leaf” categories followed the same pattern, respectively. This makes sense from a structural standpoint in that buccolingually deep incisors with relatively thick enamel caps are better suited to resisting the high forces necessary to breach hard seed coats and the chitinous exoskeletons of many insects. Conversely, the broader, thinner incisors of herbivores are better suited for shearing tough vegetation.

Of the 37 training species missed by our dietary model, only one herbivore was misclassified as a seed eater and no seed eaters were misclassified as herbivores. Thus, for all but one of the species misidentified, the misidentification was to the dietary category nearest in morphological space (i.e., omnivores fall morphologically in between herbivores and seed eaters; Fig 1). Nonetheless, it is notable that the herbivore misclassified as a seed eater, and one of the two species for which all four specimens were misclassified, was the acacia rat, *Thallomys paedulcus* (the other three specimens were misclassified as omnivores). While this species consumes the pods of acacia trees, it is reported to only consume the fleshy green outer coating and focuses the rest of its diet on young tree leaves. As such, we were compelled to categorize it as an herbivore. However, *T. paedulcus* habitually uses its incisors to cut pod-bearing twigs from acacia trees in order to carry them back to its nest for consumption [35]. It seems reasonable that a similar dental adaptation to eating hard seeds (i.e., buccolingually deep incisors with thickened enamel) would be favored for the need to cut through woody material. Therefore, as Croft et al. [28] note, because many rodents do not solely use their dentition for dietary purposes, other behaviors should be considered in discussions of dental adaptation.

### Confounding variables

The diets of many African rodents are poorly understood. One reason for this is that many rodents are true generalists that can shift dietary (and other) behaviors based on the environmental parameters of their given habitat. Murids, in particular, are known for their ecological flexibility (e.g., *Mus musculus*, the common house mouse) and over half of our specimens are members of this family. As such, the dietary information recorded for many taxa can reflect regional variations more than species-wide dietary proclivities, which we recognize.

For example, in the Kwazulu-Natal Drakensberg, *Rhabdomys pumilio*, the widely distributed and highly successful four-striped grass mouse, has a recorded diet based on 89 stomachs in which 94% contained seeds, 28% contained green plant material, and 48% contained arthropods [36]. In contrast, individuals living in Kgalagadi Transfrontier Park (Northern Cape, South Africa) had stomachs containing “93.7% herbage, 6.3% seeds, and no insects” [37].

In this regard, the diets recorded for some species, such as *Mastomys coucha*(a similarly abundant and widely distributed species), that are based on studies from a single location (22 individuals sampled in Kruger National Park by Watson [38]) must be used with caution. Of course, this does not mean that these, and other dietary data, should be disregarded, only that the inherent difficulties in accurately describing the diets of broadly distributed, generalist feeders must be considered.

We are also acutely aware of recent debates about the taxonomy of many of the rodents we have investigated (e.g., *R. pumillio* which has been separated into multiple species; see [39]). However, most, if not all of the current debates about rodent taxonomy are being resolved at the molecular level. As we are ultimately interested in applying our methods to the paleontological record, such distinctions are beyond the scope of our abilities. Simply put, the fossil record we are interested in, as of now, only provides morphological evidence.

### Pilot investigation of fossil material

In order to apply our models to fossil specimens, we chose to use random samples of rodent incisors from two sites understood to be both temporally and environmentally distinct. Isolated left maxillary rodent incisors from Swartkrans Member 1 (SKX-1) (*N*=35) and Sterkfontein Member 4 (STS-4) (*N=*45) were sampled from collections at the Ditsong Museum, South Africa, and were classified using our QDA model (Figure 3). While the sites lie roughly a kilometer away from each other on either side of the Blaaubank stream, STS-4 has been dated to ∼ 2.5 Ma [40, 41] and SKX-1 has been dated to ∼1.5 Ma [41, 43–46]. The older deposit contains *Australopithecus africanus* while the younger contains *Paranthropus robustus* and non-hominin fauna that are generally indicative of a more open environment [29, 9].

Our model predicts a strong separation in the number of specimens classified as seed eaters and herbivores in these deposits (Fig 4 and Table 3). No specimens were classified as seed eaters in the younger deposit from Swartkrans, while the older deposit from Sterkfontein contains far fewer specimens classified as herbivores (7% vs. 49%).

**Fig 4.**
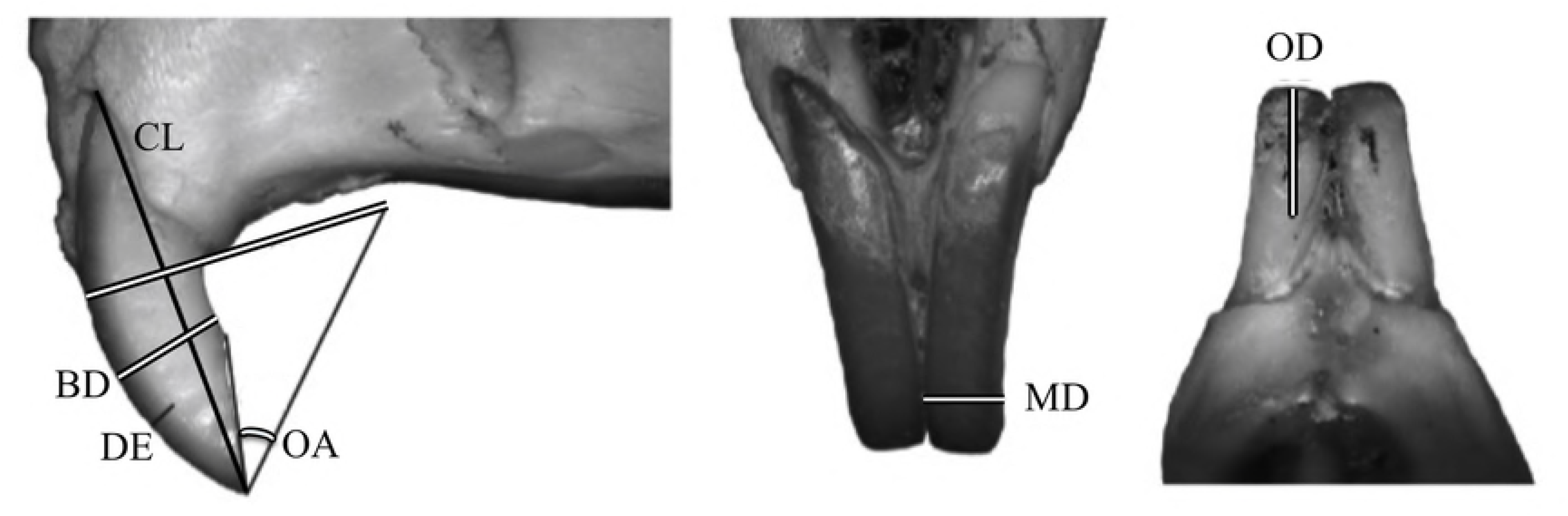
Fossils analysis. Samples from Sterkfontein Member 4 (purple circles) and Swartkrans Member 1 (orange crosses) were treated as unknowns in our dietary model. Ellipses represent 50% of the distribution of the training specimens where blue = seed eater; green = omnivore; red = herbivore.

**Table 3.** Dietary classifications of fossil specimens.

|  | **Dietary Classification** |  |  |
| --- | --- | --- | --- |
| **Fossil Site** | Herbivore | Omnivore | Seed Eater |
| SKX-1 | 17 | 19 | 0 |
| STS-4 | 3 | 26 | 16 |

SKX-1= Swartkrans Member 1; STS-4 = Sterkfontein Member 4

We are reluctant to make inferences about paleohabitat based on these results without taking taphonomic, and other potential biases in our samples and the broader deposits into account. Because such an analysis is beyond the scope of this study, we treat the fossil evidence as preliminary. We do note, however, that our sample was random and that the deposits of interest are believed to be roughly isotaphonomic [3], so it is unlikely that the comparison between these two fossil sites is of the apples to oranges variety. As such, the divergent dietary signal we reveal in our analysis might be informative about paleohabitat. In particular, none of the modern seed eater specimens we used in our model are described as living in woodland or forested environments, suggesting that our Sterkfontein sample was drawn from a more open, grassy landscape compared to our Swartkrans sample.

These results agree with previous paleoenvironmental reconstructions based on larger fauna [e.g., 9, 29] in the sense that Sterkfontein and Swartkrans differ. However, the large-bodied mammals recovered at these sites suggest that the Sterkfontein deposit accumulated during a period with less grassy cover than was found later at Swartkrans— the opposite of our conclusion [46]. More pertinent to this study, the divergent signal we find in our data does not align with Avery’s [3] taxonomic reconstructions of the two sites using small mammals, which describe both as “ecotone between grassland and savanna biomes.” Perhaps this discrepancy between our results and Avery’s [3] reflects elements of the small mammal assemblages that are under-represented (because of incompleteness) using taxonomic methods? Or, perhaps our ecomorphological signal is identifying woodland taxa that use their incisors for non-dietary behavior as seed eaters, such as *Thallomys*?

Regardless, the implications of our results are not entirely clear, as detailed studies mapping rodent diet at the community level to specific habitat parameters are lacking. Such work is a clear prerequisite before the methods used here can be fully implemented as a tool for reconstructing paleohabitats.

## Conclusion

Isolated rodent incisor morphology is a potential source of paleoenvironmental information. It may offer many of the same benefits of taxonomically-based small mammal studies, such as relatively high spatial resolution, while greatly increasing the available sample and limiting assumptions about the behavior of fossil taxa. Our model was able to successfully place modern specimens into broad dietary categories, and further refinement of our understanding of modern African rodent ecology should sharpen this model’s interpretive power.

While there appears to be a strong ecomorphological signal in isolated rodent incisors, we wish to point out that they *can* be sorted using traditional methods at broader taxonomic levels. For instance, the grooved, large, adult incisors of the otomyinae are easily recognizable as are the distinctive, stout incisors of the mole rats. Similarly, Pocock (1987) has argued that subtle grooves on the incisors can distinguish specimens belonging to the species *Mystromys albicaudatus*(an overwhelmingly dominant taxon in South African fossil assemblages that is currently listed on the IUCN Red List as endangered). As such, the future of rodent ecomorphological models using isolated incisors may be in their combined use with some degree of taxonomic sorting. However, as these ecomorphological techniques progress, it is clear that isolated fossil rodent incisors need no longer be regarded as offering limited insight into the past. Instead, they can be regarded as valuable data that paleontologists can apply to both the hominin and non-hominin fossil record.

## Acknowledgements

We wish to thank the staff at the Ditsong Museum of Natural History in Pretoria, South Africa for access to their collections. We would also like to thank the staff at the National Museum in Bloemfontein, South Africa for their help with pilot work that led to this publication. This is a research product, in whole or in part, of the Nutritional and Isotopic Ecology Lab (NIEL) at CU Boulder.

## Supporting information

**S1 Table.** Dietary categorizations and incisor measurements (mm) of rodent species sampled (OA measured in degrees). See text for explanation of dietary categories.

**S2 Table.** Blomberg’s *K* for all measures. Values > 1 suggest that phylogeny is driving the similarities between closely related species. Only values < 1 were used in our analysis.

**S3 Table.** Classification of validation specimens using our QDA model. Misclassifications are in bold. See text for explanation of dietary categories.

